# *Isoquercetin treatment of mouse sickled red blood cells shows a* discernible deformability and sickling phenotype

**DOI:** 10.64898/2026.04.24.720679

**Authors:** Osamede C. Owegie, Ivan Hancco Zirena, Tanvi Penubothu, Ionita C. Ghiran, Moua Yang

**Author notes:** Corresponding Author: Moua Yang, PhD, Bloodworks Northwest Research Institute, Division of Hematology and Oncology, Department of Medicine, University of Washington School of Medicine, 1551 Eastlake Ave E, Suite 100, Seattle, WA 98102.

## Abstract

**Introduction:** Sickle cell disease is an inherited hemoglobinopathy with defective red cell deformability. The defective deformability promotes microvascular occlusion and subsequent vaso-occlusion in sickle cell disease patients. Previous studies have demonstrated that thiol isomerases, an endoplasmic reticulum-resident oxidoreductase that is released from vascular cells into the bloodstream, are present on red cell membrane and contribute to cellular dehydration and sickling. However, the role of membrane-bound thiol isomerases on sickled red blood cells is unclear.

**Methods:** Using red blood cells from Townes humanized sickle cell or non-sickled mice, we performed ektacytometry assay under shear using laser assisted optical rotational cell analyzer (LORRCA) to assess the effects of antagonizing thiol isomerases with isoquercetin and a functional blocking monoclonal antibody. The densitometric properties of sickled red blood cells in the presence of isoquercetin was also tested using magnetic levitation.

**Results:** Thiol isomerase antagonism increased sickled red cell elongation, cellular dehydration and the diamagnetic signature compared to control treatment.

**Conclusion:** Thiol isomerases may be involved in regulating sickled red blood cells mechanical properties through mechanisms that require further investigation.

## Introduction

Sickle cell disease is an inherited hemoglobinopathy that is complicated by painful vaso-occlusive crises, a hallmark of the disease that leads to organ damage and early mortality [1]. The pathophysiology for vaso occlusion is multifaceted with red blood cell sickling and subsequent loss of membrane deformability as major contributing factors. Red cell sickling is induced by deoxygenation, promoting hemoglobin polymerization and fiber formation, which disrupts cellular morphology, increases cellular stiffness, and impacts the membrane adhesive properties. The phenotypic changes during red cell sickling can be captured by standard brightfield microscopy or through biophysical methods. Advances in ektacytometry allow for monitoring the dynamics of red cell sickling during deoxygenation using laser assisted optical rotational cell analyzer (LORRCA). The LORRCA is an ektacytometer that measures laser diffraction pattern of the red blood cells while in shear stress with a gradient of oxygen [2]. This approach can measure the point of sickling and the red cell elongation index during both deoxygenation and reoxygenation, allowing for multiparameter measurements of red cell sickling and deformability. In addition, magnetic levitation is a biophysical method that characterizes cells based on their densitometric and magnetic properties [3]. Red blood cells from various disorders, including malaria infection and sickle cell anemia, are denser and/or have increased paramagnetic properties, unlike healthy red blood cells. These properties can be captured and quantified by standard brightfield microscopy and complement many of the measurements reported by the LORRCA.

Thiol isomerases are a class of 21 oxidoreductases that catalyze the formation, cleavage, and rearrangement of disulfide bonds during protein folding [4]. Protein disulfide isomerase (PDI) is the archetypal thiol isomerase. Although it is an endoplasmic reticulum resident protein, PDI escapes ER retention, being secreted into the extracellular space, where it localizes to the surface of activated vascular cells. Vascular cells, including platelets [5, 6], endothelial cells [7], and leukocytes [8, 9], are the best characterized cell types that secrete PDI to the blood circulation. Extracellular PDI promotes blood clotting being a viable therapeutic target with galloylated polyphenols [10] and flavonoids [11]. Indeed, isoquercetin, a rutin-derived flavonoid that binds with high affinity to PDI and inhibits its reductase activity [11, 12], is currently under evaluation in clinical trials as a strategy to reduce thrombotic risk in cancer patients [13]. In addition, isoquercetin was tested in a clinical trial with sickle cell anemia patients (NCT04514510) to determine its efficacy in decreasing the risk for thrombotic complications [14]. This study also reported potential benefits in decreasing coagulation and platelet aggregation in sickle cell disease patients compared to healthy individuals [14]. Both trials found isoquercetin treatment tolerable with no reported bleeding events in the patients [13, 14]. Despite its potential role as an antithrombotic target, the mechanisms by which circulating PDI impacts other blood cells is hitherto unknown.

Mass spectrometry studies on the total red cell proteome revealed that thiol isomerases are present on red blood cells [15]. The membrane-bound PDI on red blood cells interacts with the Gardos channels (KCNN4) supporting endothelin-1 (ET-1) function during red cell sickling [16]. The presence of PDI and other thiol isomerases on red blood cells indicate a potential function for the enzymes in regulating disulfide switches that are yet to be characterized. In this study, we hypothesize that antagonizing thiol isomerases will impact the sickling phenotype and mechanical properties of red blood cells, which were investigated this using the LORRCA and magnetic levitation.

## Methods

### Reagents

Sodium citrate anticoagulant, 3.8% w/v, Hanks’ balanced salt solution (HBSS) containing calcium and magnesium with no phenol red (HGBSS^++^), and Critoseal were purchased from ThermoFisher Scientific. OxyIso was from RR Mechatronics. Townes humanized non-sickled (AA) and sickled (SS) mice were from the Jackson Laboratory. Sodium metabisulfite, dimethylsulfoxide, and isoquercetin were purchased from Sigma Aldrich. Gadoteridol Injection containing Gadolinium (Gd^+^) was from ProHance. Ketamine and xylazine were from MWI Cencora, Inc. Mouse IgG2a isotype control and anti-PDI monoclonal RL90 antibody were from ThermoFisher Scientific. N52-grade NdFeB magnets were from K&J Magnetics.

### OxyScan on the Laser assisted optical rotational cell analyzer (LORRCA)

7-12 week old Townes humanized sickle (SS) or non-sickle (AA) mice were anesthetized with 125 mg/kg ketamine and 12.5 mg/kg xylazine or with 0.2% isoflurane prior to exsanguination of whole blood into 3.8% w/v sodium citrate anticoagulant. Whole blood count was performed using a HESKA Element HT5. Red blood cells were then diluted to 2x 10^8^ into 5 mL of Oxy ISO buffer. 1.6 mL of the sample/Oxy ISO solution was injected into the LORRCA and the OxyScan program was immediately initiated. Acclimatization occurred for 2 min, deoxygenation for 21.6 min, and reoxygenation for 5.6 min. The elongation index (EI) was recorded over time during this process.

### Magnetic Levitation

Magnetic levitation was performed as previously described [3]. In short, whole blood from Townes SS or AA mice was obtained by submandibular facial vein puncture into 20 µL 3.8% w/v sodium citrate anticoagulant. The red blood cells were washed 3 times in HBSS by centrifugation at 500 x g for 10 min each. Then, two µL of the red blood cells was diluted into HBSS^++^ containing 50 mM Gd^+^ with or without 20 mM sodium metabisulfite. The red blood cell suspension was loaded into disposable 50×1×1mm square section capillary tubes by using a P200 pipette (Gilson) tip loaded with 56uL of cell suspension. During the loading and the sealing of the ends, the capillary was kept horizontally at all times. The pipette tip was was brought into close proximity to one of the ends of the capillary and the cell suspension loaded slowly into the capillary. The ends of the capillary tube were then sealed with Critoseal to prevent leakage and cells from drifting during levitation. Then capillary tube was placed into the 1 mm space between two N52-grade NdFeB magnets placed with same pole facing each other. The cells were allowed to reach levitation equilibrium for 10 minutes prior to brightfield imaging on an Olympus AX70 Provis microscope with a 20×0.40 LCPlan Fluorite objective and a condenser that provides full Kohler illumination to the sample (**Supplemental Figure**) The images were acquired with a PCO Edge 4.2 CMOS camera (Excelidas, Pittsburgh, PA) controlled by ImageJ software (NIH).

To determine the relative levitation height, the images were converted to 8 Bit in FIJI (Version: 2.16.0/1.54p) and auto-thresholded to create a binary image. The relative distribution of the cells across the image was measured as a density profile histogram across the capillary.

### *E. coli* expression of recombinant human PDI

The cDNA for human PDI was cloned into a pT7-FLAG-SBP-1 vector (ThermoFisher) between the FLAG tag and the Streptavidin Binding Peptide. The plasmid was transformed into BL21 (DE3) *E. coli* (New England Biolabs) and PDI was expressed by growing the *E. coli* in LB-ampicillin broth at 37 °C until expression was induced with 1 mM isopropyl β-D-1-thiogalactopyranoside (IPTG, CAS 367-93-1) at 24°C. The cells were lysed by sonication and the clarified lysate was poured over a high-capacity streptavidin-coated agarose column. The column was washed with 20 mM Tris-HCl pH 7.4 (CAS 1185-53-1), 150 mM NaCl (CAS 7647-14-5), 2 mM ethylenediaminetetraacetic acid (EDTA, CAS 60-00-4), and 5 mM dithiothreitol (DTT, CAS 3483-12-3). The protein was eluted with 5 mM biotin before dialyzing against phosphate buffered saline (PBS) and stored at -80°C until use.

### PDI Insulin Reductase Assay

The reductase activity of PDI was measured by a continuous spectrophotometric insulin transhydrogenation assay as previously described [17]. In brief, recombinant human PDI was diluted in a Potassium Phosphate assay buffer (61 mM K_2_HPO_4_, 39 mM KH_2_PO_4_, 2 mM EDTA, pH 7.2) to 900 nM (final reaction concentration: 450 nM) and incubated with DMSO, 50 µM isoquercetin, and 10-60 µg/mL IgG2a isotype control or anti-PDI RL90 monoclonal antibody for 30 min. 100 mM Dithiothreitol (DTT) was freshly prepared and diluted in assay buffer to 1.6 mM (final reaction concentration: 400 µM). A stock of 1.5 mM insulin was diluted to 400 µM in assay buffer to a final reaction concentration of 100 µM. The appropriate volume of DTT and insulin were combined in 200 µL of reaction in clear flat-bottom 96-well plate. Absorbance at 650 nm was recorded every minute for up to 1 hr using a Tecan Infinite Pro M Plex plate reader.

### PDI Di-eosin-GSSG reductase Assay

Recombinant PDI was diluted to 50 nM in a 100 mM potassium phosphate buffer containing 2 mM EDTA, pH 7.0. The reductase activity was monitored by fluorescence at 550 nm for up to 90 min at room temperature after the addition of 300 nM di-eosin-GSSG and 5 µM DTT.

### Statistics

The data was analyzed by two sample *t* test. A *p* value of <.05 was considered statistically significant. Data was represented as mean ± SEM.

## Results

That thiol isomerases regulate the Gardos channel through Endothelin-1 [16] and the presence of thiol isomerases in red cell proteomics [15] suggested a potential role for these oxidoreductases in regulating red cell function. To test this hypothesis, we performed ektacytometry using the LORRCA with OxyScan to profile the elongation index of sickled SS and non-sickled AA mouse red blood cells during deoxygenation and reoxygenation under shear. Baseline measurements with AA and SS red blood cells were performed. As anticipated, AA non-sickled red blood cells showed minimal changes in the elongation index throughout the deoxygenation and reoxygenation period (**Fig. 1a**). In contrast, the sickled SS red blood cells showed a characteristic decrease in EI (Elongation Index) once the point of sickling (PoS) was reached, which is measured by a 5% change in EI. The decrease in EI reaches minimal EI at 20 mmHg pO_2_. Upon reoxygenation to 120 mmHg pO_2_, the SS red blood cells undergo rapid EI recovery to baseline measurements and is similar to OxyScan profiles of human sickle red blood cells [18].

**Fig. 1.**
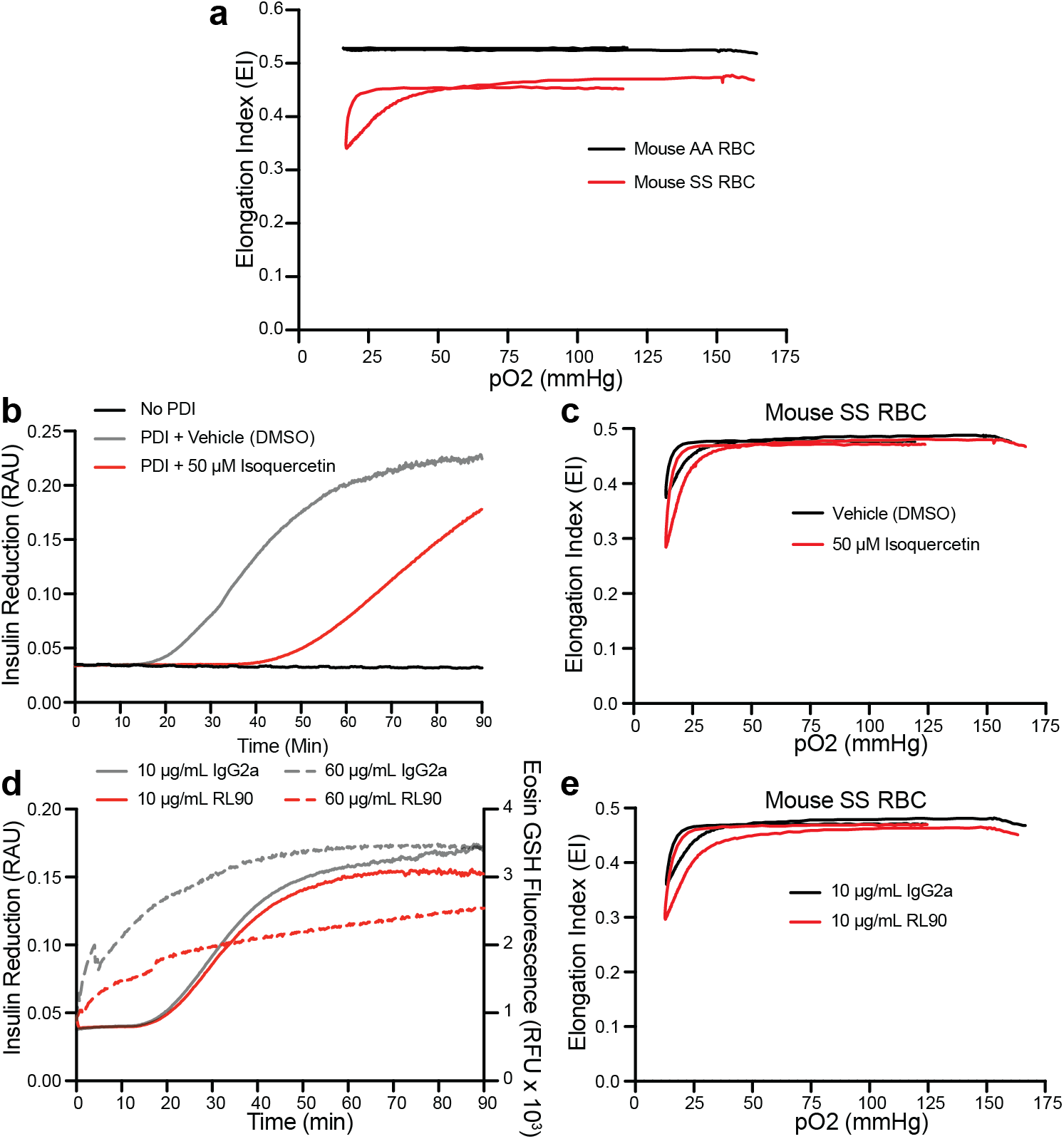
Mouse SS red blood cells show increase change in elongation index when thiol isomerase antagonist isoquercetin or RL90 were applied. **(a)** Washed red blood cells (200 x 10^6^) from AA or SS mice were diluted into Oxy Iso buffer prior to initiating the OxyScan protocol on the LORRCA. **(b)** Insulin reduction over time was measured by absorbance spectroscopy at 650 nm in the absence (black line) or presence of PDI that was treated with vehicle (DMSO, grey line) or 50 µM Isoquercetin (red line). As the disulfides on insulin are reduced by PDI, insulin precipitates over time. **(c)** Washed mouse SS red blood cells were treated with vehicle or 50 µM isoquercetin for 30 min at room temperature prior to diluting into Oxy Iso buffer for OxyScan. **(d)** Reduction of the disulfides of non-fluorescent dieosin-GSSG to fluorescent eosin-GSH by PDI was measured by spectroscopy at 550 nm (Right Y-axis; dotted lines). Insulin disulfide reduction by PDI was measured by absorbance spectroscopy at 650 nm (Left Y-axis; solid lines). **(e)** Washed mouse SS red blood cells were treated with 10 µg/mL IgG2a or RL90 for 30 min at room temperature prior to diluting into Oxy Iso buffer for OxyScan.

We treated washed mouse SS red blood cells with isoquercetin at 50 µM concentration, which is well above the reported IC_50_ value on PDI [19]. At 50 µM concentration, antagonistic effects against PDI’s disulfide breaking reductase activity were observed (**Fig. 1b**). Isoquercetin treatment consistently showed an increase in the extent of EI (delta EI), which is observed by further decrease of EI during deoxygenation compared to vehicle DMSO-treated cells (**Fig. 1c, Table 1**). There was no apparent statistical difference to the maximal EI or minimal EI with isoquercetin compared to vehicle treatment (**Table 1**). However, PoS was increased from 36 ± 3.2 mmHg pO_2_ in vehicle-treated cells to 56 ± 7.6 mmHg pO_2_ with isoquercetin treatment. These data indicate that thiol isomerases present on red blood cells, enhances SS red cell elongation and accelerates sickling. To further validate that thiol isomerases are antagonized by isoquercetin, we used the functional blocking PDI monoclonal antibody RL90, which was used to prevent PDI reductase activity from recombinant proteins, cells, and *in vivo* in a murine model of thrombus formation [5, 20]. RL90 had limited effects on PDI reductase activity at 10 µg/mL by showing a relative Vmax of 0.0039 RAU/min compared to IgG2a control antibody at 0.0045 RAU/min in the insulin reductase activity assay, an apparent ∼5-10% inhibition (**Fig. 1d, left axis**). RL90 at 60 µg/mL showed a much more robust ∼70% inhibition in the dieosin GSSG reductase activity assay with a relative Vmax of 127 RFU/min compared to IgG_2a_ of 441 RFU/min (**Fig. 1d, right axis**). These percentage of inhibition were consistent with the reported efficacy window for the RL90 antibody [5]. Nonetheless, we chose a lower concentration (10 µg/mL) due to the fact that the level of membrane-bound PDI is likely a 1-2 log-fold below this concentration since plasma PDI levels in sickle cell disease patients are in the nanomolar ranges [14]. Treating SS red blood cells with RL90 further increased the extent of EI on SS red blood cells (increase in delta EI) compared to IgG2a control antibody (**Fig. 1e, Table 1**), providing further evidence that the EI effects with isoquercetin are likely through thiol isomerase antagonism and not secondary to the chemical properties of the compound.

**Table 1.**
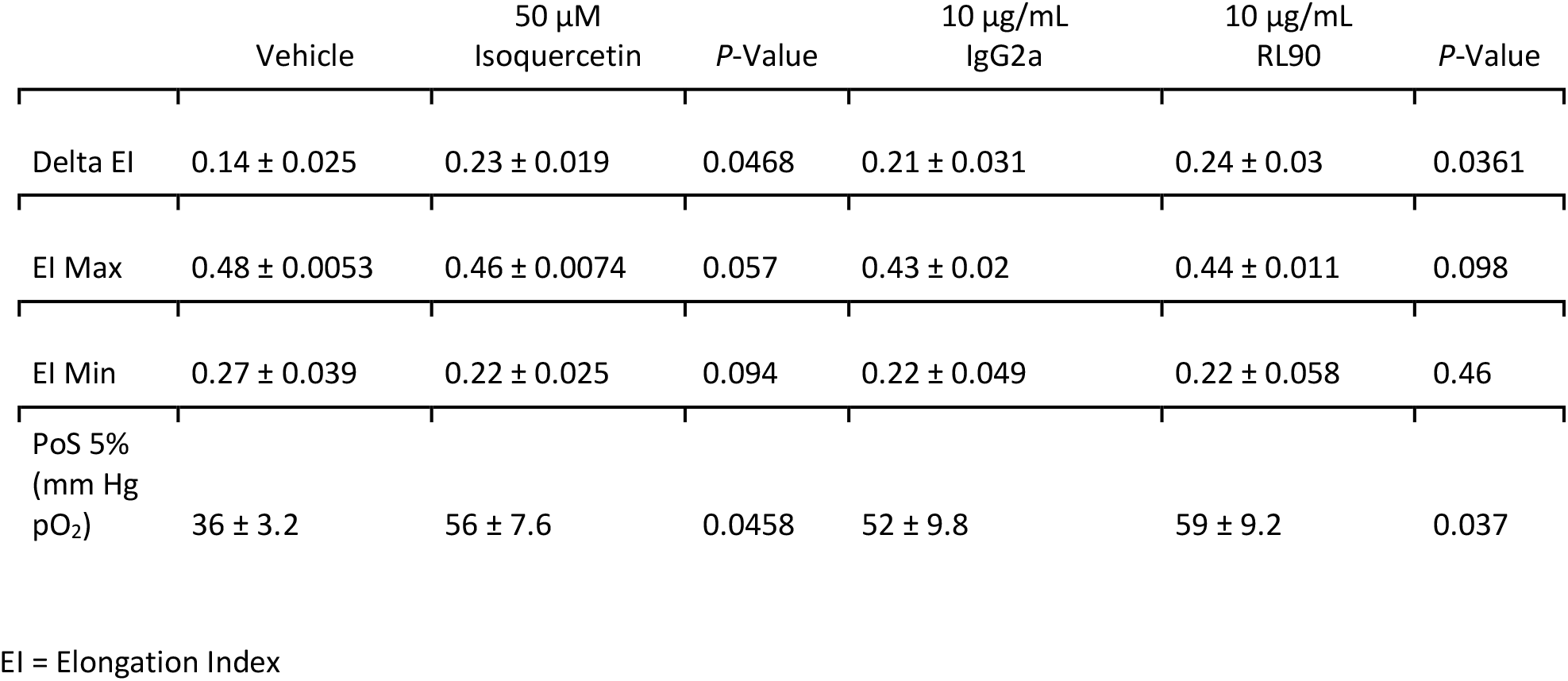
Oxyscan parameters in mouse SS RBC treated with the thiol isomerase antagonist isoquercetin or functional blocking monoclonal antibody RL90.

The increase in total EI under shear could be due to several mechanisms including hemoglobin S polymerization, cellular O_2_ tension, dehydration, and cytoskeleton dynamics [21]. Magnetic levitation can effectively test these parameters to determine sickled red cell density and magnetic properties compared to non-sickled cells in a define gadolinium-based buffer [3]. In this assay, healthy red blood cells being diagmanetic, will levitate to a height dictated by their density, whereas sickled red blood cells which are denser and more paramagnetic will move towards the bottom magnet (**Fig. 2a**) [22]. Using RBCs from a healthy human donor (AA), we treated the cells with 50 µM isoquercetin or vehicle DMSO control in the presence or absence of 15 mM dehydrating agent sodium metabisulfite (**Fig. 2b**). Isoquercetin had no effect on the RBC levitation or density in healthy human cells. We then treated SS and AA mouse red blood cells with DMSO in the presence of 20 mM sodium metabisulfite and subjected the cells to magnetic levitation. As expected, the AA mouse red blood cells were resistant to dehydrating and sickling and levitated within 10 min to a characteristic band of cells based on their density and paramagnetism (moving away from the magnet) (**Fig. 2c**). In contrast, the SS mouse red blood cells showed a curtain of cells (diamagnetism) by moving towards the magnet due to their dense nature after sickling with metabisulfite (**Fig. 2c**). Treatment of SS mouse red blood cells with isoquercetin showed an increase in cell density compared to vehicle treatment. These findings suggest that antagonizing thiol isomerases with isoquercetin may impact hemoglobin S polymerization, cellular O_2_ tension, dehydration, and cytoskeleton dynamics to induce a discernible change in SS red cell density and magnetic properties.

**Fig. 2.**
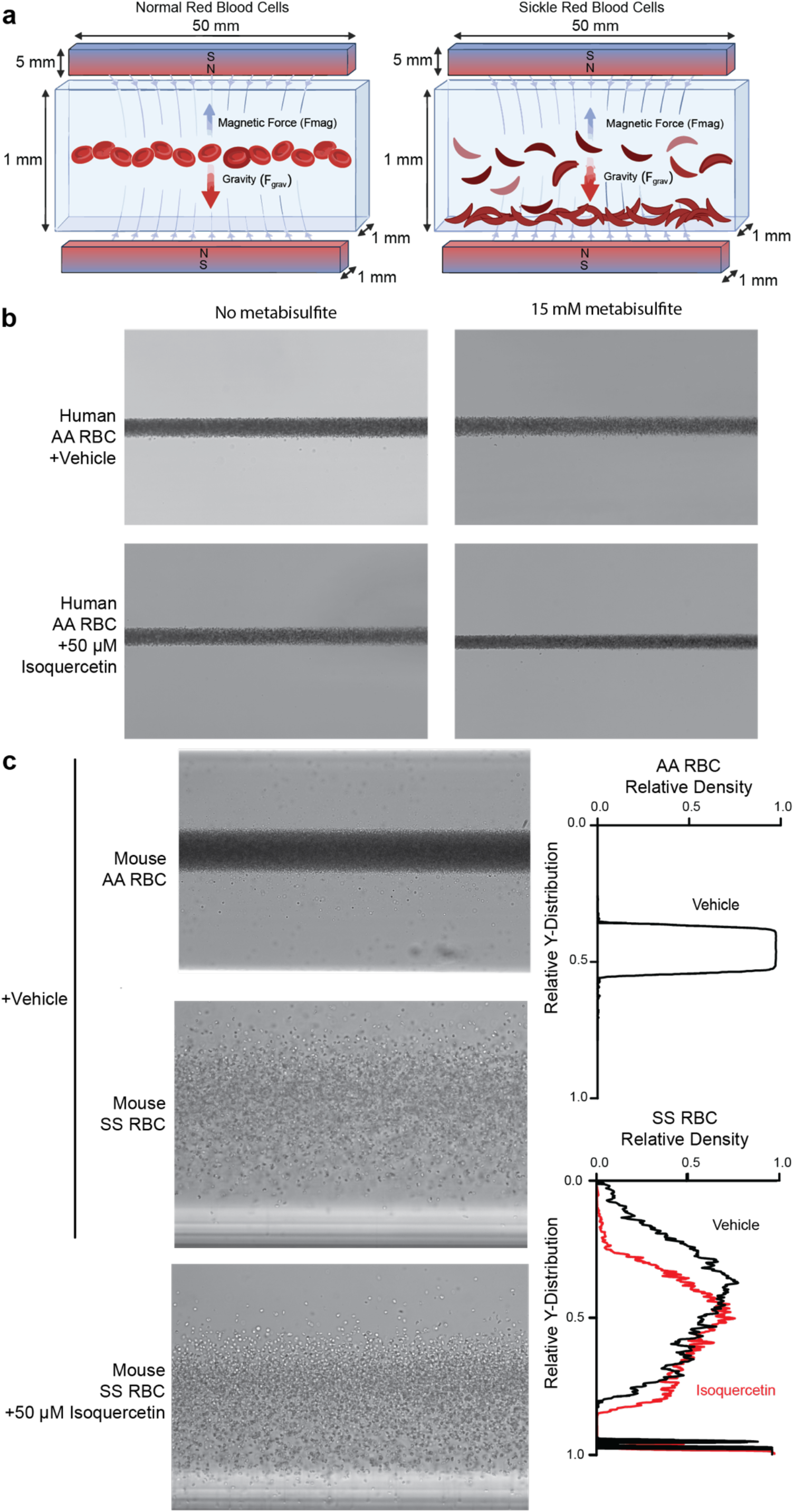
Isoquercetin treatment slightly increases mouse SS red blood cell density and diamagnetism when dehydrated with sodium metabisulfite. Red blood cells from healthy human donor, non-sickle AA Townes mice or SS sickle Townes mice were isolated and washed in HBSS containing calcium and magnesium. The cells were then treated with 50 µM Isoquercetin or vehicle control prior to resuspension in 50 mM Gd^+^ with 20 mM sodium metabisulfite or 15 mM sodium metabisulfite for mouse or human cells, respectively. The red cell suspensions were drawn into a microcapillary and placed between two N52-grade NdFeB magnets and allowed to equilibrate for 10 min before imaging. **(a)** Cartoon representation of dehydrated AA and SS RBC during magnetic levitation. SS RBCs show a curtain of dense and diamagnetic cells whereas AA RBCs are less dense and levitated to a band of cells. **(b)** Isoquercetin treatment of AA health human RBCs show no difference in levitation or density in the presence or absence of metabisulfite. **(c)** Isoquercetin treated SS mouse RBCs show a slight increase in cells levitating towards the magnet compared to vehicle treated cells, indicating a slightly elevated densit.

## Discussion

Red blood cell sickling is characteristic of sickle cell disease and is a major contributing factor to the pathophysiology of the disorder. Although red blood cells are anucleate and do not have organelles, proteomics studies have identified a plethora of proteins classically associated with various organelles [15]. Thiol isomerases are a family of 21 oxidoreductases that are highly concentrated in the endoplasmic reticulum with some of the family members secreted to the extracellular environment [4]. The extracellular thiol isomerases regulate critical disulfide bonds of membrane proteins important for thrombus formation [4]. Some of these proteins include integrins [23-25], tissue factor[26], vitronectin[27], glycoprotein Ibα[28], Factor XII [29], Histidine-Rich Glycoprotein (HRG) [30], Thrombospondin 1, Factor V[31, 32], and others [4]. Although most of these substrates are for thrombus formation, we do not know how extracellular thiol isomerases impact red cell function.

In this preliminary study we found that the flavonoid antagonist of PDI, isoquercetin, promoted red cell sickling in a shear-based ektacytometer. Specifically, the delta EI of the sickled red cell increased during deoxygenation. This increase in elongation index was also observed in sickled red blood cells treated with the monoclonal blocking RL90 antibody to PDI, providing additional evidence for the role for thiol isomerases in red cell sickling. These findings suggest that thiol isomerases may have protective function on red blood cells by regulating membrane disulfides. Although we do not know what the substrates are on the red cell membrane, prior studies have investigated the role of PDI on regulating the Gardos channel activity to decrease red cell density using the PDI antagonist, bacitracin and the RL90 blocking antibody [16]. In this former study, red cell surface-associated PDI reductase activity was increased on red blood cells from sickle SS mice compared to those from control AA mice. In the patient SS red blood cells, PDI reductase activity correlated with Gardos channel activity and PDI antagonism decreases cellular dehydration [16]. The presence of Endothelin-1 promotes the Gardos channel activity through PDI and casein kinase II-dependent mechanism [16]. Similarly, we found that antagonizing the membrane-attached PDI on SS red blood cells with isoquercetin and the blocking monoclonal antibody RL90 decreased PDI reductase activity. In contrast, treating SS red blood cells with the isoquercetin slightly enhanced red cell density and sickling. This may be related to the inhibition of the Gardos channel; however, assaying for specific Gardos channel activity is required. Although we cannot exclude the possibility of membrane-bound PDI reductase activity not captured in our assays, our findings suggest that the oxidoreductase activity and cell surface disulfide landscape in regulating red cell sickling are more complex than previously understood.

A major limitation of our study is that isoquercetin and the blocking monoclonal antibody RL90 can cross react with other thiol isomerase family members. It is unlikely that one family member of thiol isomerase contributes to red cell function, as multiple members are found in red blood cell proteomics, including PDIA3, PDIA4, and PDIA6, and TXNDC5 [15]. In addition, we previously found that many of these thiol isomerase family members are also present in the soluble fractions of red cell extracellular vesicles [33]. Nonetheless, there is a hypothesis that these family members function in an electron chain of events (e.g. donating electrons to each other through as series of cysteine reduction) based on the pKa properties of their conserved CXXC catalytic sites [34]. This hypothesis is supported by the fact that if one thiol isomerase is inhibited or genetically deleted from target cells, thrombus formation in vivo in mice induced by vessel injury is abrogated [35, 36, 20]. Both antagonists in our study showed an increase in red cell elongation, findings consistent with the hypothesis that multiple thiol isomerase family members regulate red cell sickling. Complementing these findings by identifying the disulfide landscape of the surface proteins that are either reduced or oxidized on red blood cells will be critical to understanding the mechanism of how thiol isomerases regulate red cell function.

Although PDI antagonism in SCD patients show a somewhat protective effect on coagulation and platelet aggregation potentials [14], our data suggest that on sickled mouse red blood cells PDI may regulate the sickling phenotype during deoxygenation. The net effect of this regulation with PDI’s role in platelet aggregation and coagulation will determine the therapeutic benefits of antagonism with isoquercetin. At 1000 mg/dose, which is the amount provided to SCD or pancreatic cancer patients, there were no adverse bleeding complication and no clinically adverse health effects [13, 14]. In this setting, the potential function of membrane-bound thiol isomerase on red blood cells may be more important during sickling or in hypoxic conditions when the pathogenic event (*e*.*g*. vaso-occlusive crises or clot) has already occurred. These events are also associated with oxidative stress and reactive oxygen species generation [37]. It is possible that the reactive thiols on the CGHC catalytic motif of PDI functions as scavengers of reactive oxygen species by sensing the membrane redox status of the cells. Nonetheless, these data underscores additional studies that are required to further understand how PDI regulates red cell membrane disulfides or how reactive oxygen species of SS red blood cells are influenced by PDI antagonism.

## Conclusion

We report preliminary data indicating that the flavonoid antagonist to thiol isomerases, isoquercetin, promotes a consistant decrease in mouse sickle red cell deformability and increased in cell density. Mechanistic understanding of how thiol isomerases regulate these observed phenotype will require further investigations.

## Statements

All animal studies were performed in accordance with the Institutional Animal Care and Use Committee of Bloodworks Northwest Research Institute.

## Acknowledgement

We thank Mai Shoua Vang for assisting with data visualization and John Killian Ryan for assisting with the LORRCA studies.

## Author Contributions

Conceptualization: I.C.G., M.Y.; Data Curation: O.C.O., M.Y.; Formal Analysis: O.C.O., T.P., I.C.G., M.Y.; Funding Acquisition: I.C.G., M.Y.; Investigation: O.C.O., T.P., I.H.Z., I.C.G., M.Y.; Methodology: O.C.O., I.H.Z., I.C.G., M.Y.; Project Administration: M.Y.; Resources: I.C.G., M.Y.; Supervision: M.Y.; Validation: O.C.O., I.C.G.; Visualization: M.Y.; Writing – Original Draft: M.Y.; Writing – Review & Editing: O.C.O., I.H.Z., I.C.G., M.Y.

## Data Availability Statement

The data that support the findings of this study are available from the corresponding author upon request.

