## Supplemental Figure for "*Isoquercetin treatment of mouse sickled red blood cells shows a* discernible deformability and sickling phenotype"

*Brief Report*

**Supplemental Contents**

Supplemental Figure . . . . . . . . . . . . . . . . . . . . . . . . . . . . 2

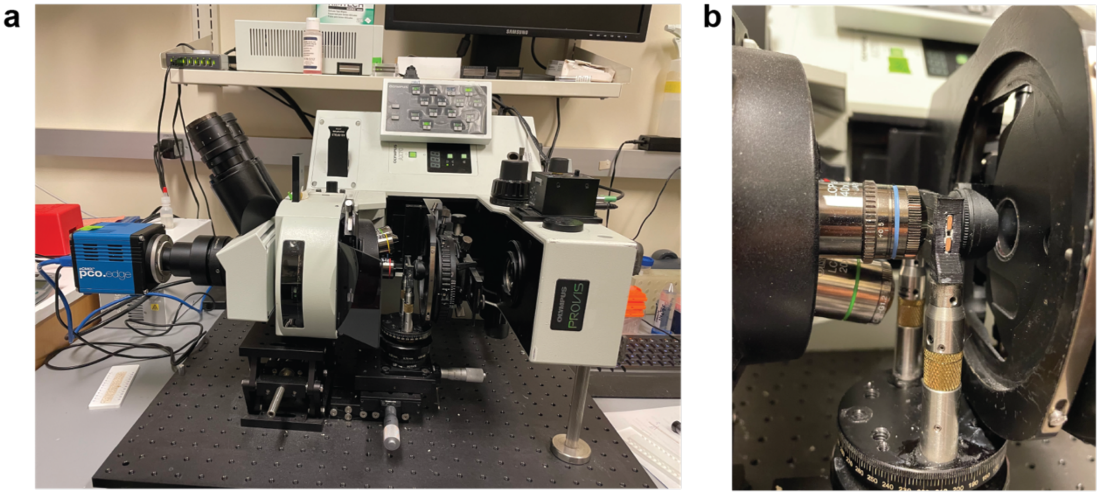


**Supplemental Figure. Magnetic levitation setup on an Olympux AX70 Provis microscope.** (A) The Olympus AX70 Provis microscope was positioned on its side and anchored and supported by Thorlab brackets onto a breadboard. (B) The two magnets were supported by a custom magnet holder held in place by two posts on a translational X,Y,Z Thorlab stage. The microscope objective is brought proximal to the capillary that is inserted between the magnets.
